# Allele Phasing Greatly Improves the Phylogenetic Utility of Ultraconserved Elements

**DOI:** 10.1101/255752

**Authors:** Tobias Andermann, Alexandre M. Fernandes, Urban Olsson, Mats Töpel, Bernard Pfeil, Bengt Oxelman, Alexandre Aleixo, Brant C. Faircloth, Alexandre Antonelli

**Author notes:** Corresponding author: Tobias Andermann, Department of Biological and Environmental Sciences, University of Gothenburg, Carl Skottsbergs Gata 22B, SE-413 19, Göteborg, Sweden.

## Abstract

Advances in high-throughput sequencing techniques now allow relatively easy and affordable sequencing of large portions of the genome, even for non-model organisms. Many phylogenetic studies reduce costs by focusing their sequencing efforts on a selected set of targeted loci, commonly enriched using sequence capture. The advantage of this approach is that it recovers a consistent set of loci, each with high sequencing depth, which leads to more confidence in the assembly of target sequences. High sequencing depth can also be used to identify phylogenetically informative allelic variation within sequenced individuals, but allele sequences are infrequently assembled in phylogenetic studies.

Instead, many scientists perform their phylogenetic analyses using contig sequences which result from the *de novo* assembly of sequencing reads into contigs containing only canonical nucleobases, and this may reduce both statistical power and phylogenetic accuracy. Here, we develop an easy-to-use pipeline to recover allele sequences from sequence capture data, and we use simulated and empirical data to demonstrate the utility of integrating these allele sequences to analyses performed under the Multispecies Coalescent (MSC) model. Our empirical analyses of Ultraconserved Element (UCE) locus data collected from the South American hummingbird genus *Topaza* demonstrate that phased allele sequences carry sufficient phylogenetic information to infer the genetic structure, lineage divergence, and biogeographic history of a genus that diversified during the last three million years. The phylogenetic results support the recognition of two species, and suggest a high rate of gene flow across large distances of rainforest habitats but rare admixture across the Amazon River. Our simulations provide evidence that analyzing allele sequences leads to more accurate estimates of tree topology and divergence times than the more common approach of using contig sequences.

---

Massive Parallel Sequencing (MPS) techniques enable time- and cost-efficient generation of DNA sequence data. Instead of using MPS to sequence complete genomes, many researchers choose to focus their sequencing efforts on a set of target loci to lower costs while achieving higher coverage and more reliable sequencing of these target regions (Faircloth et al. 2012, 2013; Mirarab et al. 2014; Smith et al. 2014; Faircloth 2015; Harvey et al. 2016; Meiklejohn et al. 2016). These multilocus datasets typically contain hundreds or thousands of target loci, and most are generated through enrichment techniques such as sequence capture (synonym: target enrichment, Gnirke et al. (2009)). After collecting sequence data from these targeted loci, many researchers assemble their high coverage sequence reads into “contigs” using *de novo* genome assembly software, and the “contig sequence” output by these assemblers often ignore the variants at heterozygous positions that are expected in diploid organisms. Typically, variable positions are treated as sequencing errors and assembly algorithms output “contig sequences” containing the more probable (i.e., numerous) variant while discarding the alternative (Iqbal et al. 2012). As a result, the “contig sequences” that are produced contain only canonical nucleobases, losing the information about read variability at variable positions. Hereafter, we use “contigs” and “contig sequences” to refer to the sequences that are output by *de novo* assemblers.

One alternative approach to generating contig sequences uses the depth of sequencing coverage to programatically identify variable positions within a targeted locus (also known as “calling” single nucleotide polymorphisms (SNPs)) and subsequently sorting (or “phasing”) these SNPs into two allele sequences or “haplotypes” which represent alleles on the same chromosome present at that locus. These approaches have been used to estimate demographic parameters such as effective population size, rate of migration, and the amount of gene flow between and within populations. However, it is rarely acknowledged (*c.f.* Lischer et al. 2014; Potts et al. 2014; Schrempf et al. 2016; Eriksson et al. 2017) that allelic sequences are useful for phylogenetic studies to improve the estimation of gene trees, species trees, and divergence times (Garrick et al. 2010; Potts et al. 2014; Lischer et al. 2014). The common practice of neglecting allelic information in phylogenetic studies possibly results from historical inertia and a lack of computational pipelines to prepare allele sequences for phylogenetic analysis using MPS data.

In addition to the problems of determining allelic sequences, the proper analysis of allelic information in phylogenetic studies remains a challenging and intensively discussed topic (Garrick et al. 2010; Lischer et al. 2014; Potts et al. 2014; Schrempf et al. 2016; Leach´e and Oaks 2017). Various approaches have been proposed to include this information into phylogenetic methods (Lischer et al. 2014; Potts et al. 2014; Schrempf et al. 2016). One is to code heterozygous sites using the International Union of Pure and Applied Chemistry (IUPAC) ambiguity codes and to include these as additional characters in existing substitution models for gene tree and species tree inference (Potts et al. 2014; Schrempf et al. 2016). While these studies demonstrate that integrating additional allelic information in this manner increases accuracy in phylogenetic inference, Lischer et al. (2014) found that coding heterozygous sites as IUPAC ambiguity codes in phylogenetic models biases the results toward older divergence time estimates. Instead, Lischer et al. (2014) introduced a method of repeated random haplotype sampling (RRHS) in which allele sequences are repeatedly concatenated across many loci, using a random haplotype for any given locus in each replicate. In their approach, they then analyzed thousands of concatenation replicates separately for phylogenetic tree estimation and summarized the results between replicates, thereby integrating the allelic information in the form of uncertainty intervals. However, there are two important shortcomings of this approach: 1. concatenating unlinked loci (and in particular allele sequences from unlinked loci) in a random manner is known to produce incorrect topologies (Degnan and Rosenberg 2009) often with false confidence (Edwards et al. 2007; Kolaczkowski and Thornton 2004; Kubatko and Degnan 2007; Mossel and Vigoda 2005), which is not accounted for when doing so repeatedly and summarizing the resulting trees, and 2. running thousands of tree estimation replicates based on extensive amounts of sequence data results in unfeasibly long computation times, particularly for Markov-Chain Monte Carlo (MCMC) based softwares such as MrBayes or BEAST. Hence, there is need to find proper solutions to include heterozygous information in phylogenetic analyses, as concluded by Lischer et al. (2014).

Here, we introduce the bioinformatic assembly of allele sequences from UCE data (Fig. 1) and demonstrate a full integration of allele sequences to species tree estimation under the multispecies coalescent (MSC) model. In our approach, we treat each allelic sequence of an individual at a given locus as an independent sample from the population, and we analyze these sequences using the species tree and delimitation software STACEY (Jones et al. 2014; Jones 2017), which allows for this approach by not requiring *a priori* clade- or species-assignments. We first demonstrate the empirical utility of this approach by resolving the shallow genetic structure (<1 Ma) within two recognized morphospecies of the South American hummingbird genus *Topaza*, with a dataset of 2,386 ultraconserved elements (UCEs, see Faircloth et al. (2012)). We then validate this approach, using simulated data, and we find evidence that allele sequences yield more accurate results in terms of species tree estimation and species delimitation than the contig sequence approach that ignores heterozygous information. Further, our simulation results provide evidence that compiling phased allele sequences and treating these as individual samples outperforms alternative approaches of coding heterozygous information, such as analyzing sequences containing IUPAC ambiguity codes or analyzing isolated SNPs. We conclude that allele phasing for sequence capture data can be critical for correct species delimitation and phylogeny estimation, particularly in recently diverged groups, and that analyses using phased allele sequences should be considered as one, potential “best practice” for analyzing sequence capture datasets in a phylogenetic context.

**Figure 1:**
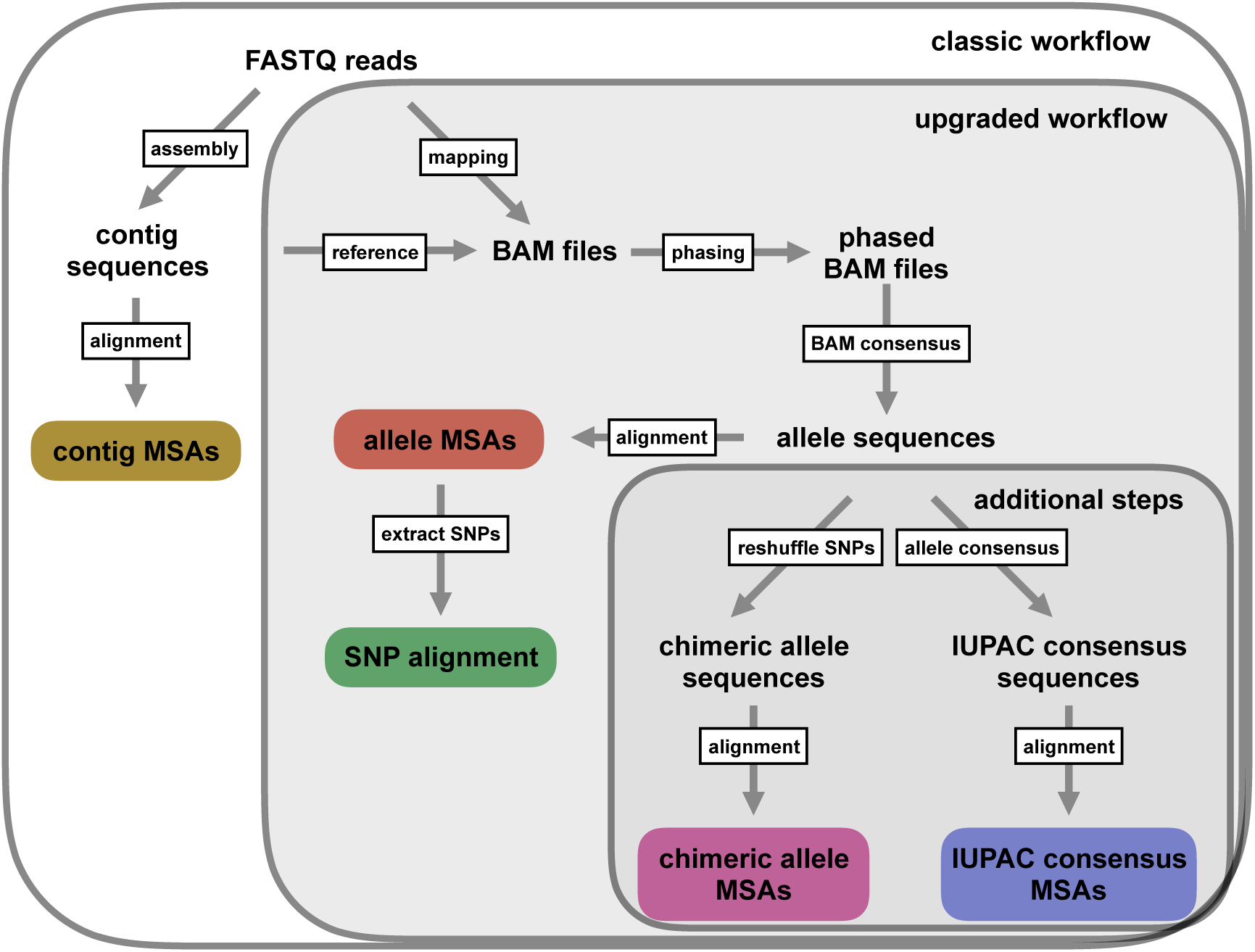
Depiction of the workflow used in this manuscript. Colored boxes represent different types of multiple sequence alignments (MSAs) used for phylogenetic inference in this study. In addition to the standard UCE workflow (boxlabel: classic workflow) of generating contig MSAs (Faircloth et al. 2012; Smith et al. 2014; Faircloth 2015), we extended the bioinformatic processing in order to generate UCE allele MSAs, and to extract single nucleotide polymorphism (SNPs) from these allele MSAs (boxlabel: upgraded workflow). We added these new functions to the PHYLUCE pipeline (Faircloth 2015). Additional data processing steps (boxlabel: additional steps) were executed in this study in order to test different codings of heterozygous positions.

## Materials and Methods

### Study System

The genus *Topaza* and its sister genus *Florisuga* form the Topazes group, which together with the Hermits represent the most ancient branch within the hummingbird family (Trochilidae) (McGuire et al. 2014). Topazes are estimated to have diverged as a separate lineage from all other hummingbirds around 21.5 Ma, whereas the most recent common ancestor (MRCA) of *Topaza* and *Florisuga* lived approximately 19 Ma (McGuire et al. 2014). At present, there are two morphospecies recognized within *Topaza*, namely the Fiery Topaz, *T. pyra* (Gould, 1846), and the Crimson Topaz, *T. pella* (Linnaeus, 1758). However, the species status of *T. pyra* has been challenged by some authors (Schuchmann 1999; Orn´es-Schmitz and Schuchmann 2011), who consider this genus to be monotypic. Topaz hummingbirds are endemic to the Amazonian rainforest and are some of the most spectacular and largest hummingbirds worldwide, measuring up to 23 cm (adult males, including tail feathers) and weighing up to 12 g (Schuchmann et al. 2016; del Hoyo et al. 2016a). These birds are usually found in the forest canopy along forest edges and clearings, and are often seen close to river banks (Orn´es-Schmitz and Schuchmann 2011). There is morphological evidence for several subspecies within both currently recognized *Topaza* species (Peters 1945; Schuchmann 1999; Hu et al. 2000; Ornés-Schmitz and Schuchmann 2011) that we investigate using genetic data.

### Sequence Data Generation

We extracted DNA from the muscle tissue of 10 vouchered hummingbirds (9 *Topaza*, one *Florisuga*, see Table 1) using the Qiagen DNeasy Blood and Tissue Kit according to the manufacturer’s instructions (Qiagen GmbH, Hilden, Germany). These samples cover most of the genus’ total geographic range (Fig. 2) and all morphologically recognized intraspecific taxa (Schuchmann et al. 2016; del Hoyo et al. 2016a). All samples were sonicated with a Covaris S220 to a fragment length of 800 base pairs (bp). Paired-end, size-selected (range 600-800bp) DNA libraries were prepared for sequencing, using the magnetic-bead based NEXTflexTM Rapid DNA-Seq Kit (Bioo Scientific Corporation, Austin, TX, USA), following the user’s manual (v14.02).

**Figure 2:**
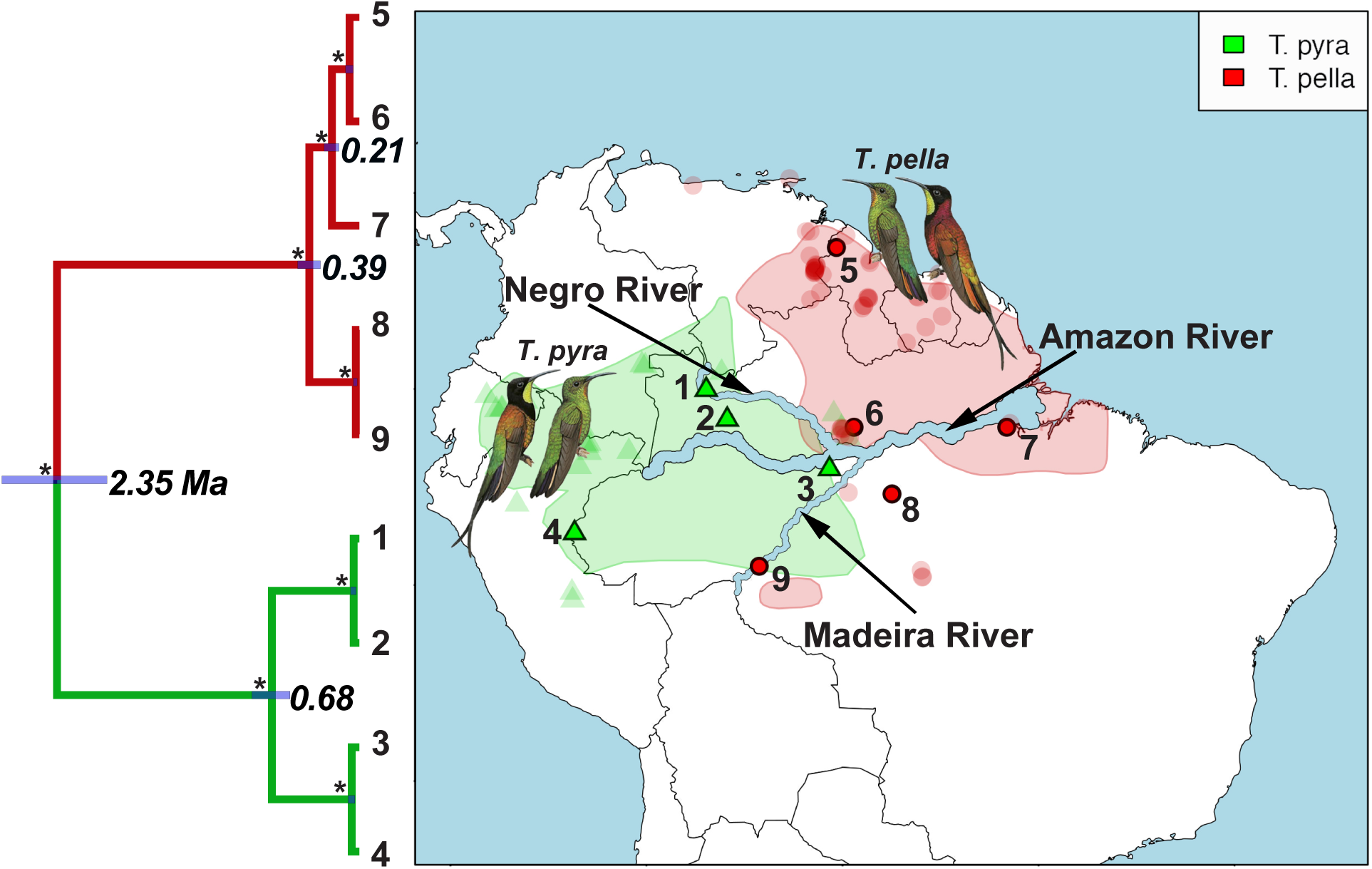
Distribution ranges and mitochondrial phylogeny of the South American hummingbird genus *Topaza*. Tip labels of phylogeny and numbers on map represent sample IDs (Table 1) of sequenced *Topaza* specimens. Node labels in phylogeny show mean divergence time estimates for mitochondrial lineages, with node bars representing the surrounding uncertainty (95% highest posterior density (HPD)). All nodes are supported with 100% posterior probability (PP), as indicated by asterisks. Polygons on map represent distribution ranges of the two morphospecies (*T. pyra* and *T. pella*) as estimated by BirdLife International (http://www.birdlife.org). Transparent symbols (triangles and circles) represent *Topaza* sightings, which were downloaded from the eBird database (Sullivan et al. 2009). The major river systems in the Amazon drainage basin are labeled and emphasized in size for better visibility. *Topaza* illustrations were provided by del Hoyo et al. (2016b).

**Table 1.** Sequenced specimens and coordinates of their sampling locations, subspecies identifications based on morphological characters, abbreviation for sample providers: INPA = Instituto Nacional de Pesquisas da Amazoˆnia, MPEG = Museum Paraense Em´ılio Goeldi, USNM = NMNH, Smithsonian Institution, Washington DC, USA.

| ID | Taxon | Subspecies | Voucher number | Latitude | Longitude |
| --- | --- | --- | --- | --- | --- |
| 1 | *Topaza pyra* | *amaruni* | INPA A1106 | -0.044167 | -66.94944 |
| 2 | *T. pyra* | *pyra* | MPEG 62475 | -1.559444 | -65.88006 |
| 3 | *T. pyra* | *pyra* | MPEG 62474 | -4.083889 | -60.66050 |
| 4 | *T. pyra* | *pyra* | MPEG 52721 | -7.350000 | -73.66667 |
| 5 | *T. pella* | NA | USNM 586322 | 7.220000 | -60.29000 |
| 6 | *T. pella* | *pella* | INPA A3319 | -1.927900 | -59.41600 |
| 7 | *T. pella* | *smaragdula* | MPEG 61688 | -1.950000 | -51.60000 |
| 8 | *T. pella* | *microrhyncha* | MPEG 65603 | -5.352417 | -57.47500 |
| 9 | *T. pella* | NA | INPA A6233 | -9.028550 | -64.24231 |
| 10 | *Florisuga fusca* | NA | MPEG 70697 | -15.15972 | -39.04500 |

We used the “Tetrapods-UCE-2.5Kv1” bait set (uce-2.5k-probes.fasta), consisting of 2,560 baits (each 120 bp), targeting 2,386 UCEs, as described by Faircloth et al. (2012). The bait sequences were downloaded from http://ultraconserved.org and synthesized by MYcroarray (Biodiscovery LLC, Ann Arbor, MI, USA). Sequence enrichment was performed using a MYbaits kit according to the enclosed user manual (v1.3.8). The enriched libraries were then sequenced using 250 bp, paired-end sequencing on an Illumina MiSeq machine (Illumina Inc., San Diego, CA, USA). Library preparation, sequence enrichment and sequencing were performed by the Sahlgrenska Genomics Core Facility in Gothenburg, Sweden.

### Mitochondrial Genome

To infer a dated mitochondrial phylogeny for the genus *Topaza* to compare with the nuclear phylogeny, we used off-target mitochondrial reads to assemble the complete mitochondrial genome for all samples. We found that as many as 4.5% of all sequence reads were of mitochondrial origin, even though no baits targeting mitochondrial loci were used during sequence capture. An alignment of the assembled mitochondrial genomes for all samples was analyzed in BEAST (Drummond et al. 2012). Dating priors included clock-rate priors for three mitochondrial genes, estimated for honeycreepers by Lerner et al. (2011) and node-age priors within the genus *Topaza* that were estimated by McGuire et al. (2014). The resulting phylogeny and estimated divergence times are shown in 2. A detailed description of the assembly and phylogenetic analysis of the mitochondrial genome data can be found in online Appendix 1 (Supplemental Material available on Dryad, doi:10.5061/dryad.hq3vq).

### UCE Data Processing

For this study we generated five different types of datasets, which we analyzed under the MSC. These five datasets represent different coding schemes for heterozygous information and are listed and described in the following sections.

#### 1. UCE contig alignments

Because contig sequences are commonly used in phylogenetic analyses of MPS datasets (e.g. Faircloth et al. (2012); Smith et al. (2014); Faircloth (2015)), we generated multiple sequence alignments (MSAs) of contigs for all UCE loci in order to test the accuracy of the phylogenetic estimation of this approach.

To create MSAs from UCE contig data, we followed the suggested workflow from the PHYLUCE documentation (http://phyluce.readthedocs.io/en/latest/tutorial-one.html). We applied the PHYLUCE default settings unless otherwise stated. First we quality-filtered and cleaned raw Illumina reads of adapter contamination with Trimmomatic (Bolger et al. 2014), which is implemented in the PHYLUCE function illumiprocessor. The reads were then assembled into contigs using the software ABYSS (Simpson et al. 2009) as implemented in the PHYLUCE pipeline. In order to identify contigs representing UCE loci, all assembled contigs were mapped against the UCE reference sequences from the bait sequence file (uce-2.5k-probes.fasta), using the PHYLUCE function match_contigs_to_probes.py. We extracted only those sequences that matched UCE loci and that were present in all samples (n=820). These UCE sequences were then aligned for each locus (Fig. 1) using MAFFT (Katoh et al. 2009).

#### 2. UCE allele alignments

We altered the typical UCE workflow in order to retrieve the allelic information that is lost when collapsing multiple reads into a single contig sequence (Fig. 1). To create this new workflow, we extracted all UCE contigs for each sample separately and treated each resulting contig set as a sample-specific reference library for read mapping (reference-based assembly). We then mapped the cleaned reads against each reference library on a per sample basis, using CLC-mapper from the CLC Workbench software. The mapped reads were sorted and then phased with SAMtools v0.1.19 (Li et al. 2009), using the commands samtools sort and samtools phase, respectively. This phasing function is based on a dynamic programming algorithm that uses read connectivity across multiple variable sites to determine the two phases of any given diploid locus (He et al. 2010). Further, this algorithm uses paired-end read information to reach connectivity over longer distances and it minimizes the problem of accidentally phasing a sequencing error, by applying the minimum error correction function (He et al. 2010).

UCE data provide an excellent dataset for allele phasing based on read connectivity, because the read coverage across any given UCE locus typically is highest in the center and decreases toward the ends. This makes it possible to phase throughout the complete locus without any breaks in the sequence. Even in cases where the only variable sites are found on opposite ends of the locus, the insert size we targeted in this study (800 bp), in combination with paired-end sequencing, enabled the phasing process to bridge the complete locus (average length of compiled UCE-sequences in our study was 870 bp).

The two phased output files (BAM format) were inspected for proper variant separation for all loci using Tablet (Milne et al. 2013). We then collapsed each allele BAM file into a single consensus sequence per haplotype and exported the two resulting allele sequences for each sample in FASTA format. In order to separate true heterozygous sites from occasional variants introduced by sequencing errors, we only made a nucleotide call if the respective nucleotide was supported by at least three reads. Ambiguous positions were coded with the IUPAC code ‘N’ in the allele consensus sequences. We explored the difference in the treatment of heterozygous positions between the contigs produced by the *de novo* assembler ABYSS and our phased allele sequences in detail (exemplary for one sample) in online Appendix 2 (Supplemental Material).

In the next, step we aligned the allele sequences between all samples, separately for each UCE locus, using MAFFT (Fig. 1). We integrated this complete workflow into the UCE processing software PHYLUCE (Faircloth 2015) with slight alterations, one of which is the use of the open-source mapping program bwa (Li and Durbin 2010) in place of CLC-mapper.

#### 3. UCE IUPAC consensus sequence alignments

We generated an additional set of alignments by merging the two allele sequences for each individual into one consensus sequence with heterozygous sites coded as IUPAC ambiguity codes (merge_allele_sequences_ambiguity_codes.py, available from: github.com/tobiashofmann88/UCE-data-management/). We used this dataset to test whether our allele phasing approach improved phylogenetic inference when compared to the IUPAC consensus approach applied in other studies, where heterozygous positions are coded as IUPAC ambiguity codes in a consensus sequence for each locus and individual (Potts et al. 2014; Schrempf et al. 2016).

#### 4. UCE chimeric allele alignments

To investigate whether correct phasing of heterozygous sites is essential or if similar results are achieved by randomly placing variants in either allele sequence, we generated a dataset with chimeric allele sequence alignments. We created these alignments by applying a custom python script (shuffle_snps_in_allele_alignments.py, available from: github.com/tobiashofmann88/UCE-data-management/) to the phased allele sequence alignments and randomly shuffling the two variants at each polymorphic position between the two allele sequences for each individual. This process leads, in many cases, to an incorrect combination of variants on each allele sequence, thereby creating chimeric allele sequences. The resulting alignments contain the same number of sequences as the phased allele alignments (two sequences per individual), whereas the contig alignments and the IUPAC consensus alignments contain only half as many sequences (one sequence per individual).

#### 5. UCE SNP alignment

A common approach to analyzing heterozygous information is to reduce the sequence information to only a single variant SNP per locus. This data-reduction approach is often chosen because multilocus datasets of the size generated in this study can be incompatible with Bayesian MSC methods applied to the full sequence data, due to extremely long computational times and convergence issues. Instead, alignments of unlinked SNPs can be used to infer species trees and species demographics under the MSC model with the BEAST2 package SNAPP (Bryant et al. 2012), a program specifically designed for such data. However, extracting and filtering SNPs from BAM files with existing software (such as the Genome Analysis Toolkit (GATK), McKenna et al. (2010)) and converting these into a SNAPP compatible format can be cumbersome, because SNAPP requires positions with exactly two different states, coded in the following manner: individual homozygous for the original state = “0”, heterozygous = “1”, and homozygous for the derived state = “2”.

To alleviate this problem, we developed a python function that extracts biallelic SNPs directly from allele sequence MSAs (snps_from_uce_alignments.py, available from: github.com/tobiashofmann88/snp extraction from alignments/). Extracting SNPs from MSAs in this manner is a straightforward and simple way to generate a SNP dataset compatible with SNAPP, and does not require re-visiting the BAM files. A similar program is also available in the R-package phrynomics (Leach´e et al. 2015). We used this approach to extract one variable position per alignment (to ensure unlinked SNPs) that had exactly two different states among all *Topaza* samples, not allowing for positions with missing data or ambiguities. This produced a SNP dataset of 598 unlinked SNPs.

### Generation of Simulated UCE Data

To assess the accuracy of the phylogenetic inferences resulting from different data processing approaches, we simulated UCE data similar to those discussed in the five processing schemes we applied to the empirical *Topaza* data. However, because this approach required us to simulate allele alignments before generating contig alignments, steps one and two, below, are reversed from their order, above. We repeated all steps involving the generation and analyses of simulated data to produce 10 independent simulation replicates.

#### 1. Simulated allele alignments

In order to simulate allele alignments similar to our empirical data we first estimated species divergence times and population sizes from the empirical UCE allele MSAs under the MSC model (Rannala and Yang 2003) using the Bayesian MCMC program BPP v3.1 (Yang 2015). We applied the A00 model, which estimates divergence times and population sizes from MSAs for a given species tree topology. As input topology we used the species tree topology resulting from the analysis of the empirical allele MSAs in STACEY, assigning the *Topaza* samples to five separate taxa (corresponding to colored clades in Figure 3b). An initial BPP analysis did not converge in reasonable computational time, a problem that has previously been reported for UCE datasets containing several hundred loci (Giarla and Esselstyn 2015). To avoid this issue, we split the 820 UCE alignments randomly into 10 subsets of equal size (n=82) and analyzed these separately with identical settings in BPP. The MCMC was set for 150,000 generations (burn-in 50,000), sampling every 10 generations. We summarized the estimates for population sizes and divergence times across all 10 individual runs. We then applied the mean values of these estimates to the species tree topology, by using the estimated divergence times as branch lengths and estimated population sizes as node values, resulting in the species tree in Figure 4g. This tree was used to simulate sequence alignments with the MCcoal simulator, which is integrated into BPP. Equivalent to the empirical data, we simulated sequence data for five taxa (D, E, X, Y, and Z) and one outgroup taxon (F, not shown in Figure 4g). In the simulations, these taxa were simulated as true species under the MSC model. In order to mimic the empirical allele data, we simulated four individuals for species ‘D’ (equivalent to two allele sequences for 2 samples), four for species ‘E’, four for species ‘X’, two for species ‘Y’ (two allele sequences for one sample), four for species ‘Z’, and two for the outgroup species ‘F’. In this manner we simulated 820 UCE allele MSAs of 848 bp length (a value equal to the average alignment length of the empirical allele alignments). The resulting simulated allele MSAs are equivalent to our empirical allele MSAs, containing two phased allele sequences for every individual that differ only in true heterozygous sites and which are not expected to contain read-errors.

#### 2. Simulated contig alignments

To simulate UCE contig MSAs that contain sequences similar to contigs generated by assemblers like ABYSS, Velvet or Trinity, which pick only one of the two variants at a heterozygous site, we merged the sequences within each coalescent species in pairs of two (equivalent to pairs of allele sequences). Each pair of allele sequences was joined into one contig sequence by randomly picking one of the two variants at each heterozygous site across all loci. As in the empirical contig assembly approach, our simulation approach may generate chimeric contig sequences.

#### 3. Simulated IUPAC consensus alignments

Next, we generated IUPAC consensus MSAs in the same manner as we generated the simulated contig MSAs in the previous step, with the exception that all heterozygous sites were coded with IUPAC ambiguity codes instead of randomly picking one of the two variants.

#### 4. Simulated chimeric allele alignments

We generated chimeric allele sequence MSAs from the simulated allele MSAs by randomly shuffling the heterozygous sites between each pair of sequences using the same pairs as in the previous two steps.

#### 5. Simulated SNP alignment

Finally, we extracted two different SNP datasets from the simulated phased allele MSAs. The first SNP dataset (SNPs complete) was extracted in the same manner as described for the empirical data (one SNP per locus for all loci) which resulted in a total alignment length of 820 SNPs for the simulated data. We extracted an additional SNP dataset (SNPs reduced) from only the subset of the 150 simulated allele alignments that were used for the sequence-based MSC analyses (see next section below). The resulting dataset of 150 SNPs was used to compare the phylogenetic inference based on SNP data versus that based on full sequence data, if the same number of loci is being analyzed. This enabled us to evaluate the direct effect of reducing the full sequence information in the MSAs to one single SNP for each of the selected 150 loci.

### MSC Analyses of Empirical and Simulated UCE Data

#### Sequence-based tree estimation

To jointly infer gene trees and species trees, we analyzed each of the generated sets of MSAs (processing schemes 1-4 for empirical and simulated) under the MSC model, using the DISSECT method (Jones et al. 2014) implemented in STACEY (Jones 2017), which is available as a BEAST2 (Bouckaert et al. 2014) package. STACEY allows *BEAST analyses without prior taxonomic assignments, searching the tree space while simultaneously collapsing very shallow clades in the species tree (controlled by the parameter collapseHeight). This collapsing avoids a common violation of the MSC model that occurs when samples belonging to the same coalescent species are assigned to separate taxa in *BEAST. This feature makes STACEY suitable for analyzing allele sequences, because they do not have to be constrained to belong to the same taxon and can be treated as independent samples from a population. STACEY runs with the usual *BEAST operators, but integrates out the population size parameter and has new MCMC proposal distributions to more efficiently sample the species tree, which decreases the time until convergence. In order to reach even faster convergence, we reduced the number of loci for this analysis by selecting the 150 allele MSAs with the most parsimony informative sites. This selection was made for both the empirical and the simulated allele MSAs. The same 150 loci were selected for all other processing schemes.

**Figure 3:**
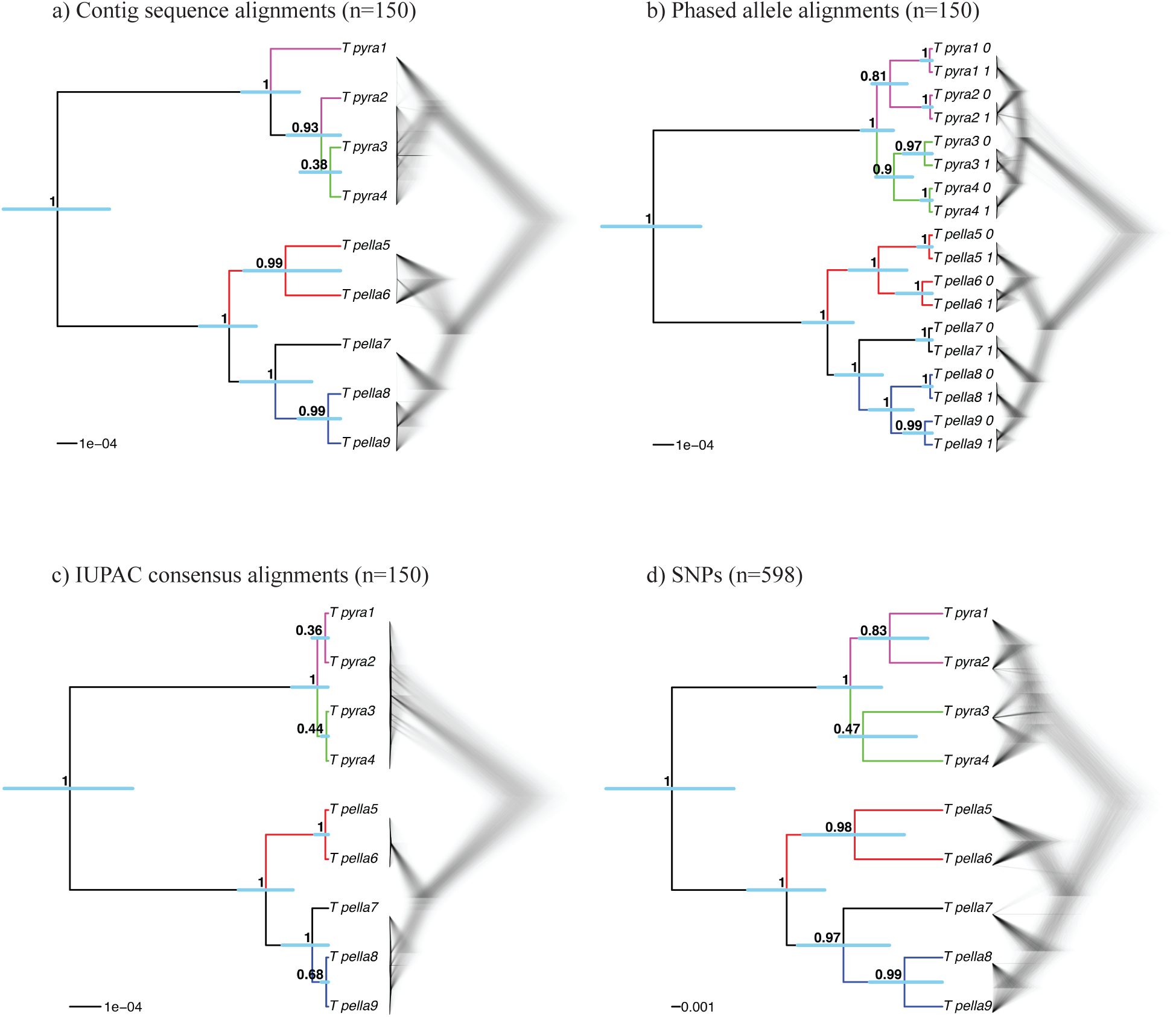
Multispecies Coalescent (MSC) species trees for the empirical *Topaza* data, based on four different data types used in this study: contig sequence MSAs, phased allele sequence MSAs, IUPAC consensus sequence MSAs and SNP data. a) STACEY species tree from UCE contig alignments (n=150), b) STACEY species tree from UCE allele alignments (n=150), STACEY species tree from UCE IUPAC consensus alignments (n=150) and d) SNAPP species tree from UCE SNP data (1 SNP per locus if present, n=598). Shown are the Maximum Clade Credibility trees (node values = PP, error-bars = 95% HPD of divergence times) and a plot of the complete posterior species tree distribution (excluding burn-in).

**Figure 4:**
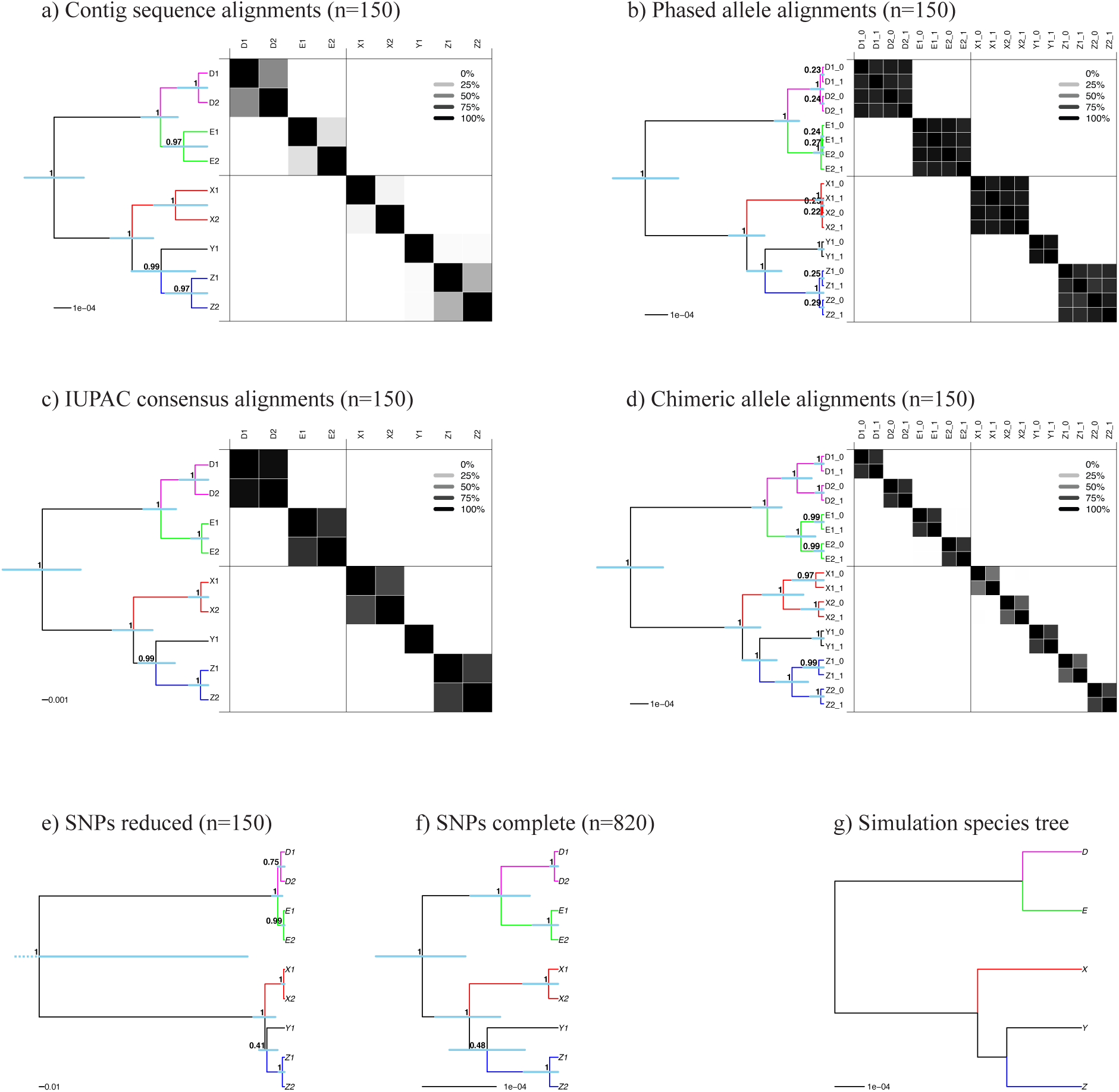
MSC species tree results for different data processing schemes of simulated data. a) to d) show the STACEY results of the four different types of MSAs analyzed in this study. Displayed in these panels are the Maximum Clade Credibility trees and the similarity matrices depicting the posterior probability of two samples belonging to the same clade, as calculated with SpeciesDelimitationAnalyser. Dark panels depict a high pairwise similarity, whereas light panels depict low similarity scores (see legend). e) and f) show the Maximum Clade Credibility trees resulting from SNAPP for our two SNP datasets, (reduced and complete). g) shows the species tree under which the sequence data were simulated in this study. Node support values in PP, blue bars representing 95% HPD confidence intervals.

Prior to analysis, we estimated the most appropriate substitution model for each of the 150 loci with jModeltest (Supplementary Table S1) using BIC. We used BEAUTI v2.4.4 to create an input file for STACEY in which we unlinked substitution models, clock models and gene trees for all loci. We did not apply any taxon assignments, thereby treating every sequence as a separate taxon. We chose a strict clock for all loci and fixed the average clock rate for one random locus to 1.0, while estimating all other clock rates in relation to this locus. To ensure that all resulting species trees were scaled to an average clock rate of 1.0, we rescaled every species tree from the posterior distribution (post analysis) using the average clock rate of the respective MCMC step. We applied the STACEY-specific BirthDeathCollapse model as a species tree prior, choosing a value of 1e-5 for the collapseHeight parameter. Other settings were: bdcGrowthRate = log normal (M=4.6, S=1.5); collapseWeight = beta (alpha=2, beta=2); popPriorScale = log normal (M=-7, S=2); relativeDeathRate = beta (alpha=1.0, beta=1.0). For the IUPAC consensus data, we enabled the processing of ambiguous sites by adding useAmbiguities=“true” to the gene tree likelihood priors for all loci in the STACEY XML file. All analyses were run for 1,000,000,000 MCMC generations or until convergence (ESS values *>*200), logging every 20,000 generations. Convergence was assessed using Tracer v1.6 (Rambaut et al. 2013). We then summarized the posterior tree distribution into one Maximum Clade Credibility tree (i.e. tree in the posterior sample that has the maximum product of posterior clade probabilities) with TreeAnnotator v2.4.4, discarding the first 10% of trees as burn-in.

For the simulated data, we analyzed the posterior species tree distributions of each analysis with the program SpeciesDelimitationAnalyser (part of the STACEY distribution). This program produces a similarity matrix that contains the posterior probabilities of belonging to the same cluster for each pair of sequences. This analysis was run with a collapseHeight value of 1e-5 (identical to the collapseHeight used in the STACEY analysis), while discarding the first 10% of trees as burn-in.

#### 2. SNP-based tree estimation

To estimate the species tree phylogeny from the extracted SNP data, we analyzed the empirical and simulated SNP data in SNAPP. We did not apply prior clade assignments to the samples in the SNP alignment (each sample was assigned as its own taxon). We set coalescent rate and mutation rates to be estimated based on the input data, and we chose a Yule species tree model with default settings (λ = 0.00765). We ran the analysis for 10,000,000 generations, sampling trees and other parameters from the posterior every 1,000 generations. Unlike STACEY, SNAPP assumes correct assignments of all sequences to coalescent species. Using the simulated SNP data, we therefore tested how our approach of assigning every individual as its own coalescent species affects the resulting phylogenetic inference. We did so by running a separate analysis for both simulated SNP datasets (complete and reduced) with correct species assignments (assignments as in Figure 4g).

### Additional Analyses

We ran additional analyses of the contig and the phased allele MSAs for both the empirical and simulated data using a summary coalescent approach as implemented in MP-EST (Yu et al. 2007), which can be found in online Appendix 3 (Supplemental Material) and Supplementary Figures S1-S3.

## Results

### UCE Summary Statistics

#### Alignment statistics

In the following we use the term “polymorphic sites” for those positions within a MSA alignment of a given locus where we find at least two different states at a particular position among the sequences for all samples. This does not require a particular individual being heterozygous for the given position, since we do not search for SNPs on a per sample basis but rather for SNPs within the genus *Topaza*. In this manner, we found that the empirical UCE contig sequence alignments had an average of 2.8 polymorphic sites per locus and an average alignment length of 870 bp. In contrast, phasing the empirical UCE data to create allele alignments led to 4.5 polymorphic sites per locus and an average alignment length of 848 bp, representing a 60% increase in polymorphic sites per locus. This increase of polymorphic sites was attributable to the fact that many variants get lost during contig assembly, because ABYSS and other tested contig assemblers, namely Trinity and Velvet, often eliminate one of the two variants at heterozygous positions (see below). The reduced length of the allele alignments in comparison to the contig alignments was due to conservative alignment clipping thresholds implemented in PHYLUCE, which clips alignment ends if less than 50% of sequences are present. Because the allele phasing algorithm divides the FASTQ reads into two allele bins and because a nucleotide is only called if it is supported by at least three high-quality FASTQ reads, we lost some of the nucleotide calls at areas of low read coverage (mostly at the ends of a locus) when comparing the allele sequences to the contig sequences. More information about the distribution of lengths and variable sites within the empirical UCE data can be found in the Supplementary Figures S4 and S5. The simulated contig MSAs had an average of 3.2 polymorphic sites per locus, after excluding the outgroup (average calculated across all 10 simulation replicates). The simulated allele MSAs, on the other hand, contained an average of 5.4 polymorphic sites (69% increase) across 10 independent simulation replicates. An overview of parsimony informative sites, variable sites and length of each alignment (simulated and empirical data) can be found in Supplementary Table S2.

### MSC Results of Empirical UCE Data

The MSC species tree results for all tested processing schemes of the empirical UCE data (contig sequences, allele sequences, IUPAC consensus sequences, chimeric allele sequences and SNPs) strongly support the monophyly of both *T. pyra* and *T. pella* with 100% Bayesian posterior probability (PP) (Fig. 3 and Supplementary Fig. S6). In all MSC analyses, we also see strongly supported genetic structure within *T. pella* (≥ 97% PP), separating the northern samples (5 and 6, sampled north of the Amazon River) from the southern ones (7, 8 and 9, sampled south of the Amazon River). Additionally, within the shallow southern *T. pella* clade, all datasets, with exception of the IUPAC consensus data (Fig. 3c), strongly support a genetic distinction (≥ 99% PP) between sample 7 from the Amazon River delta and the other southern *T. pella* samples (8 and 9). Further, the analysis of the phased allele MSAs returns a phylogenetic signal, possibly also tracking a genetic divergence between a northern and a southern clade within *T. pyra*, but their monophyly is not very strongly supported (Fig. 3b). This pattern is further supported by the mitochondrial phylogeny, which shows the same divergence within *T. pyra*, dated at 0.68 million years ago (Fig. 2 and online Appendix 1).

### MSC Results of Simulated Data

#### Species tree topology

We analyzed six different datasets under the MSC model for each of the ten simulation replicates: contig sequence MSAs (n=150, STACEY), allele sequence MSAs (n=150, STACEY), IUPAC consensus MSAs (n=150, STACEY), chimeric allele MSAs (n=150, STACEY), reduced SNP data (n=150, SNAPP), and the complete SNP dataset (n=820, SNAPP). All resulting species trees (Fig. 4a-f) correctly return the topology of the species tree that was used to simulate the data (Fig. 4g) across all ten simulation replicates (Supplementary Fig. S7). All central nodes in the species trees are supported by ≥90% PP in all analyses, with the exception of the species tree resulting from the reduced SNP dataset, which shows very weak support for two nodes and has a large uncertainty interval around the root-height (Fig. 4e). However, these shortcomings disappeared when we added more (unlinked) SNPs to the dataset (Fig. 4f). The full SNP dataset (n=820) produced the correct species tree topology with high node support consistently throughout all ten independently simulated datasets (Supplementary Fig. S8). The SNAPP species tree topology appeared to be unaffected by the chosen clade assignment model; while we allowed every sequence to be its own taxon in Figure 4e and f, we also applied the correct species assignment (as in Fig. 4g) in two additional analyses for one of the simulation replicates (reduced and complete SNP data) that returned the same tree topology (Supplementary Figs. S9 and S10).

#### Species delimitation

Although the inferred species tree topology was consistent among all four sequence-based MSC analyses (Fig. 4a-d), the inferred node heights varied considerably between the species trees resulting from the different data processing schemes. For the contig sequence data (Fig. 4a) and the chimeric allele data (Fig. 4d), the node heights within the five simulated species (D,E,X,Y,Z) were too high, which led to an overestimation of the number of coalescent species in the dataset (see similarity matrices). Conversely, the phased allele data (Fig. 4b) and the IUPAC consensus data (Fig. 4c) correctly delimited the five coalescent species from the simulation input tree (Fig. 4g). The STACEY results showed the same pattern in all ten simulation replicates (Fig. S7).

#### Accuracy of divergence time estimation

For all four sequence-based analyses (Fig. 4a-d) the average substitution rate across all loci was set to ‘1’. Under these settings, we expected the absolute values of the sequence-based analyses to return the node height values of the simulation input tree, which used substitution rates scaled in the same manner. The phased allele MSAs produced the most accurate estimation of divergence times out of all tested datasets (see proximity of estimates to simulation input value, represented by green line in Figure 5). This was the case for all nodes in the species tree, namely (D,E), (Y,Z), (X,(Y,Z)), and ((D,E)(X,(Y,Z))). The divergence time estimates resulting from the phased allele data accurately recovered the true values and did not show any bias throughout ten simulation replicates (Supplementary Fig. S11). This contrasts with the contig MSAs and the chimeric allele MSAs that consistently overestimated the height of all nodes and the IUPAC consensus MSAs which consistently underestimated the height of all nodes (Figs. 5 and S11).

**Figure 5:**
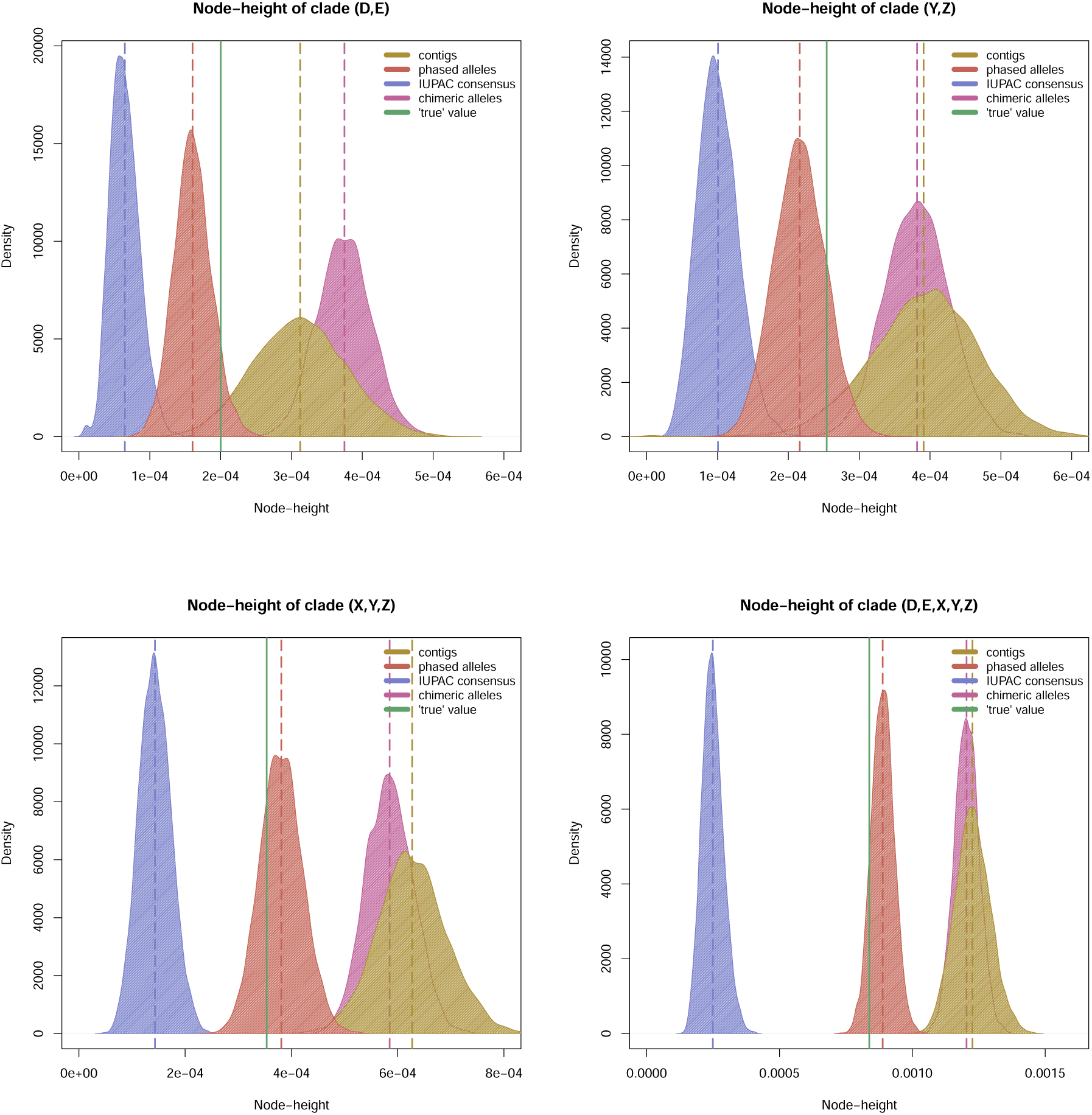
Posterior distributions of divergence times, estimated with STACEY. Each panel represents a different node in the STACEY species tree (see panel titles) and shows density plots of the posterior node-height distribution (excl. 10% burnin) for each of the 4 sequence-based processing schemes: contig sequences, phased allele sequences, IUPAC consensus sequences and chimeric allele sequences (see legend for color-codes). The dotted vertical lines show the means of these posterior distributions. The solid vertical line shows the true node height value, which is the node height for the respective clade in the input species tree, under which the sequence alignments were simulated.

## Discussion

### Phased Allele Sequences Return The Most Accurate Phylogeny

We tested whether phylogenetic inference improves by phasing sequence capture data into allele sequences, in comparison to the standard workflow of analyzing contig sequences (Faircloth et al. 2012; McCormack et al. 2012; Smith et al. 2014; Faircloth 2015). The answer is yes. We find that phased allele data outperform contig sequences in terms of species delimitation (Fig. 4) and divergence time estimation (Fig. 5). Contig sequence MSAs on the other hand lead to a consistent overestimation of divergence times (Fig. 5), which in turn lead to an overestimation of the number of coalescent species in our simulated data (Fig. 4a). These results support earlier work by Lischer et al. (2014), who concluded that consensus sequences introduce a bias towards older node heights. Because both our empirical and simulated data represent rather shallow phylogenetic relationships, future research is required to determine if these findings also apply to datasets representing divergence events occurring in deeper time.

Besides these practical advantages of using phased allele sequences for phylogenetic analyses, there are further theoretical arguments for compiling and analyzing allele sequence MSAs from sequence capture datasets.

First, allele sequences represent the smallest evolutionary unit on which selection and other evolutionary processes act. Therefore, the coalescent models that underlie our phylogenetic methods, including the MSC model Degnan and Rosenberg (2009), have been developed for allele sequences. Contig sequences, on the other hand, represent an artificial and possibly chimeric sequence construct that arises from merging all read variation at a given locus into a single sequence. This process masks information by eliminating one of the two variants at a heterozygous site (online Appendix 2). This shortcoming of the most common assemblers (e.g. ABYSS, Trinity and Velvet) is due to the fact that they were designed to assemble sequences of haploid genomes and they are not optimized for heterozygous sequences or genomes (Bodily et al. 2015).

Second, not only are allele sequences the more appropriate data type, but phasing sequence capture data also leads to a doubling of the effective sample size, since two sequences are compiled for a diploid individual, in contrast to the single sequence per individual that is recovered when taking the contig approach. Here, we demonstrate how these sequences can be properly applied as independent samples from a population by using the assignment-free BirthDeathCollapse model as implemented in STACEY. Because STACEY requires no *a priori* assignment of sequences to taxa, it avoids a violation of the MSC that would occur when analyzing allele sequences as separate taxa in *BEAST, since *BEAST assumes each taxon constitutes a separate coalescent species.

Third, sequence capture datasets such as UCEs are optimal for allele phasing because they contain high read coverage collected across short genomic intervals that are optimal for read-connectivity based phasing. The workflow developed in this study is now fully integrated into the PHYLUCE pipeline, making allele phasing for sequence capture data easily available to a broad user group.

### Phasing of Heterozygous Sites Matters

Several studies have accounted for heterozygosity by inserting IUPAC ambiguity codes into their sequences at variable positions (Potts et al. 2014; Schrempf et al. 2016), rather than phasing SNPs to produce separate allele sequences. Here, we directly compared these two approaches, and found that the IUPAC consensus sequences performed equally well to the phased allele sequences for estimating the species tree topology (Fig. 4). However, IUPAC consensus sequence data led to a consistent underestimation of the divergence times of all nodes in the species tree (Fig. 5). Our results contrast with those of (Lischer et al. 2014), who reported an overestimation of divergence times for alignments containing IUPAC ambiguity codes. The differences between our results may simply be caused by the different tree inference programs used. Lischer et al. (2014) applied a Neighbour Joining (NJ) tree algorithm as implemented in the software PHYLIP (Felsenstein 2005) that treats two sequences containing the same ambiguity codes as identical. In effect, the approach used by Lischer et al. (2014) did not directly investigate the effect of IUPAC ambiguity codes on phylogenetic estimates but rather the effect of removing heterozygous sites. Our approach of analyzing IUPAC consensus sequences under the MSC in STACEY, on the other hand, properly integrates these IUPAC ambiguity codes into the calculation of the gene tree likelihoods. Thus, we conclude that IUPAC ambiguity codes introduce a bias towards younger divergence times, even when properly integrated into the phylogenetic model. The underlying cause of this discrepancy should be further investigated in future studies.

We also tested whether the improved performance of phased allele sequences in comparison to contig or IUPAC consensus sequence data may merely be an effect of doubling the number of sequences in the MSAs, by analyzing a dataset of chimeric allele sequences with randomly shuffled SNPs. As with the contig data, the chimeric allele data led to an overestimation of the number of coalescent species (Fig. 4d) and to a biased estimation towards older divergence times (Fig. 5). The fact that contig sequences and chimeric allele sequences produce very similar results in our analyses is not surprising, because contigs, themselves, represent chimeric consensus sequences of the variation found at a locus within an individual. The similarity of the results between contig MSAs and chimeric allele MSAs also shows that the number of sequences being analyzed does not affect the estimated topology, species delimitation or divergence time estimates (Figs. 4 and 5).

Based on the findings discussed above, we conclude that proper phasing of heterozygous positions is preferable to the alternative of coding heterozygous sites as IUPAC ambiguity codes, particularly when the estimation of divergence times is of interest. Further, allele sequences are theoretically more appropriate input for coalescent models and should be the preferred data type input to these models. The scalability of this approach to larger sample sizes and the applicability of our results to studies of older divergences are questions that should be investigated in future studies.

One additional issue that we do not address in this study are the effects of sequencing errors. While sequencing errors can potentially be a serious issue particularly for datasets affected by low read coverage, we do not expect sequencing errors to be assembled into our final allele sequences, due to our relatively high read coverage per exported variant (*>*three reads each). The effects of sequencing errors and incorrectly inferred read variability on downstream analyses are subjects that need to be explored in future studies.

### Practicality of Using Phased Allele Data in Multilocus Phylogenetics

In this study, we analyze MSAs resulting from the different processing schemes in a MSC framework using the STACEY BirthDeathCollapse tree model. However, due to the size (number of samples and loci) of many sequence capture datasets, it is often unfeasible to analyze all MSAs jointly in one MSC analysis because of computational limitations (Smith et al. 2014; Manthey et al. 2016). This problem is exacerbated when working with allele MSAs compared to the contig or IUPAC consensus approach, because each alignment contains twice the number of sequences, leading to a doubling of tips in all estimated gene trees. Here we outline three different strategies of addressing this problem:

1. One reasonable approach to data reduction is to use a subset of the allele MSAs for phylogeny estimation. We chose this approach here and reduced the UCE dataset from 820 MSAs to 150 MSAs in order to reach convergence of the MCMC (BirthDeathCollapse without taxon-assignments) within a reasonable time frame (three to four days, single core on a Mac Pro, Late 2013, 3.5 GHz 6-Core Intel Xeon E5 processor). This approach has the advantage that we can fully integrate the allelic sequence information and avoid *a priori* assignments of allele sequences to taxa. However this approach discards the majority of the multilocus information by excluding most MSAs from the analysis.
2. An alternative approach to data reduction, while keeping the multilocus information of all loci, is to analyze only a single polymorphic position (SNP) per MSA using SNAPP (Bryant et al. 2012). We find that phased allele MSAs provide an excellent template for SNP extraction; since all polymorphisms present in the allele sequences have already undergone quality and coverage filters, it is very straightforward to extract SNPs directly from the allele MSAs. We provide an open-source script for this purpose which also converts the extracted SNPs into a SNAPP compatible format. In our study, this approach produced the correct species tree topology and also estimated the relative node-heights correctly (Fig. 4f). However, SNAPP can only estimate relative and not absolute values for divergence times (Bryant et al. 2012), in contrast to sequence-based analyses (Fig. 4a-d) that deliver absolute divergence time estimates. A more thorough discussion about extracting SNPs from sequence capture data can be found in online Appendix 4 (Supplemental Material).
3. Another common approach is to abdicate the more appropriate but computationally heavy co-estimation of gene trees and species trees of the MCMC-based MSC methods and chose species tree methods that separate gene tree and species tree estimation into two consecutive steps. This family of methods is often referred to as summary coalescent methods. In this approach gene trees are estimated separately for each MSA. In a subsequent step, the estimated gene trees are used to infer the most likely species tree. The advantage of this approach is that the number of independent loci being analyzed does not constitute a serious computational limitation, because every gene tree is estimated independently, which allows for efficient computational parallelization. On the other hand, summary coalescent methods are sensitive to the number of informative sites per individual locus (Gatesy and Springer 2014; Springer and Gatesy 2014). Given that our phased allele MSAs contained on average 60% more polymorphic sites than the contig MSAs (69% for the simulated data), we argue that phased allele MSAs may lead to more precise phylogenetic estimates under the summary coalescent approach in comparison to contig MSAs. In our case, the summary coalescent approach was not very suitable, due to rather conserved alignments with limited number of informative sites for individual gene tree inference, which obscured the inference of branch lengths in the species tree (online Appendix 3). However, in the case of our simulated data, we observed a more precise estimate of the species tree topology based on phased allele MSAs when compared to those based on contig MSAs (online Appendix 3). In conclusion the summary coalescent approach can be suitable if the individual alignments contain a sufficient number of parsimony informative sites for gene tree inference, and for this reason it is likely that phased allele MSAs might return more precise phylogenetic estimates than contig MSAs. However, further simulation studies are required to properly test this hypothesis.

### Phylogenetic relationships in Topaza

#### One or two species?

Our results show a separation of two lineages within the genus *Topaza* that is dated at ca. 2.4 Ma in the mitochondrial tree (Fig. 2 and online Appendix 1). These lineages are consistent with the previously described morphospecies *T. pyra* (Gould, 1846) and *T. pella* (Linnaeus, 1758) that are generally accepted in the ornithological community (Hu et al. 2000; del Hoyo et al. 2016a). However, the species status of *T. pyra* has been challenged by some authors (Orn´es-Schmitz and Schuchmann 2011; Schuchmann 1999). These authors concluded that *Topaza* is a monotypic genus with *pyra* being a subspecies of *T. pella*, which they refer to as *T. pella pyra*. Our results consistently support *T. pyra* as a separate lineage across all analyses, lending no support for the conspecificity of these two taxa (Fig. 3).

#### Genetic divergence within morphospecies

One aim of this study was to evaluate the genetic structure within the two morphospecies, *T. pyra* and *T. pella*. The mitochondrial tree shows two divergent clades within *T. pyra* (Fig. 2 and online Appendix 1), but these clades are not strongly supported by the UCE data (Fig. 3), even though the allele sequence data are picking up a signal that possibly indicates two clades are in the process of diversifying (Fig. 3b). For *T. pella*, on the other hand, we consistently find the same clades throughout all multilocus MSC analyses (Fig. 3), leading us to distinguish between the following populations that are congruent with previous morphological subspecies descriptions: a northern *T. pella* population (*T. pella pella*), a southern *T. pella* population (*T. pella microrhyncha*) and a separate population occupying the estuary region of Amazon River (*T. pella smaragdula*). We discuss these phylogenetic conclusions in more detail in online Appendix 5 (Supplemental Material).

#### Summarizing biogeographic remarks

The presence of genetically similar individuals sampled at great geographic distances (e.g. samples 5 and 6) suggests that *Topaza* hummingbirds maintain high levels of gene flow across vast distances of rainforest habitat. At the same time, we find indicators of phylogenetic structure within species, distinguishing samples that are separated by only a small geographic distance (see e.g. samples 6 and 8). These samples are however separated by the Amazon River, which has been found to constitute a dispersal barrier for various species of birds and many other animals (Remsen and Parker 1983; Clair 2003; Hayes and Sewlal 2004; Moore et al. 2008; Fernandes et al. 2012; Ribas et al. 2012; Thom and Aleixo 2015). Even though some hummingbird species are known to disperse across large distances (Wyman et al. 2004; Russell et al. 1994), the Amazon River and its associated habitats (such as seasonally flooded forests) may be part of a complex network of factors that inhibit gene flow among populations of *Topaza* hummingbirds.

## Conclusions

This study provides evidence that the assembly of phased allele sequence MSAs improves phylogenetic inference under the MSC model. We find that contig sequences, on the other hand, which are commonly used for phylogenetic inference, lead to biases in the estimation of divergence times. Additionally, phased allele sequence MSAs provide a useful template for the extraction of SNP data, and SNP data can be applied as an alternative dataset for phylogenetic inference, circumventing some computational limitations when analyzing multilocus full-sequence data with MCMC-based MSC methods. Our empirical results suggest the separation of two species within the genus *Topaza*, and we further find genetic structure within one of these species, justifying the definition of separate subspecies. Based on our empirical and simulated results, we conclude that allele phasing should be considered as one “best practice” for processing sequence capture data, although the sample-size, phylogenetic scale, and analytical limitations of this approach have not yet been well-established.

## Supplementary Material

Supplementary material, including Supplemental Figs. S1-S11, Supplemental Tables S1 and S2, online Appendices 1-5 as well as data files, can be found in the Dryad data repository at https://doi.org/10.5061/dryad.hq3vq.

## Availability

The documentation for the allele phasing workflow, which we included into the PHYLUCE pipeline, can be found here: http://phyluce.readthedocs.io/en/latest/tutorial-two.html. The script for extracting SNPs from MSAs is available here: https://github.com/tobiashofmann88/snp_extraction_from_alignments. All processing and analyses steps executed on the data are stored in bash-scripts on our project GitHub page at https://github.com/tobiashofmann88/topaza_uce. The raw sequencing reads are stored in the NCBI Short Read Archive (SRA) at https://www.ncbi.nlm.nih.gov/sra/SRP135707.

## Acknowledgments

We wish to thank all those ornithologists who have dedicated their time to collecting samples in Amazonia; museum curators for providing us with samples for this study; Brazilian authorities for issuing the permits needed for this work; our lab engineer Anna Ansebo for laboratory assistance; Alexander Zizka for assistance in creating the range maps; HBW Alive for providing the *Topaza* illustrations; and colleagues at our labs for discussions and feedback. We further thank Susanne Renner, Adam Leache and 4 anonymous reviewers for their feedback on earlier versions of this manuscript. Computational analyses were performed on the bioinformatics computer cluster Albiorix at the Department of Biological and Environmental Sciences, University of Gothenburg.

